# Towards a species search engine: KISSE offers a rigorous statistical framework for bone collagen tandem mass spectrometry data

**DOI:** 10.1101/2025.03.24.641820

**Authors:** Hassan Gharibi, Amir Ata Saei, Alexey L. Chernobrovkin, Susanna L. Lundstrom, Hezheng Lyu, Zhaowei Meng, Akos Vegvari, Massimilliano Gaetani, Roman A. Zubarev

**Affiliations:** Division of Chemistry I, Department of Medical Biochemistry and Biophysics, Karolinska Institutet, Stockholm 171 77, Sweden; Chemical Proteomics, Swedish National Infrastructure for Biological Mass Spectrometry (BioMS), Stockholm, Sweden; Chemical Proteomics Unit, SciLifeLab, Stockholm 171 77, Sweden; Department of Microbiology, Tumor and Cell Biology, Karolinska Institutet, Stockholm 171 77, Sweden; Pelago Bioscience, Solna 171 65, Sweden

## Abstract

DNA and bone collagen are two key sources of resilient molecular markers used to identify species from their remains. Collagen is more stable than DNA, and thus it is preferred for ancient and degraded samples. Current mass spectrometry-based collagen sequencing approaches are empirical and lack a rigorous statistical framework. Based on the well-developed approaches to protein identification in shotgun proteomics, we introduce a first approximation of the species search engine (SSE). Our SSE named KISSE is based on a species-specific library of collagenous peptides that uses both peptide sequences and their relative abundances. The developed statistical model can identify the species and the probability of correct identification, as well as determine the likelihood of the analyzed species not being in the library. We discuss the advantages and limitations of the proposed approach and the possibility of extending it to other tissues.

---

The identification of animal species by a piece of bone, including fossilized samples, is of utmost importance in several scientific disciplines, such as paleontology, archaeology, and forensics. This information helps scientists better understand the evolution, behavior, and interaction of the species with the environment[1]. In forensics, the identification of bone samples can aid in criminal investigations and the documentation of human remains[2]. To a large extent, the ability to accurately identify animal species from pieces of bones can advance our understanding of the evolution of the natural world and human history[3].

While genetic analysis of DNA extracted from the bone provides the most definite species ID, collagen, a protein found in connective tissues such as bones and cartilage, has been found to be more resistant to degradation and easier to extract from fossils and ancient remains than DNA[4]. This is because collagen is much more abundant in the bone than DNA, and its triple protein helix is more stable than the DNA double helix against various environmental stressors, such as heat, moisture, and exposure to ultraviolet radiation[5]. Also, collagen sequences in different tissues and organs can vary, unlike DNA, and thus offer the possibility to identify, at least to some degree, not only the species but also the tissue[6,7]. In addition, racemization and deamidation of amino acid residues in collagen can provide valuable information about the age and conditions of bone preservation, which can help researchers better understand the context of the remains[8]. Therefore, in some areas of science, collagen analysis has gained more popularity than DNA sequencing [9,10].

At first glance, the identification of species by collagen analysis should be a rather straightforward task. Collagen can be easily extracted from any bone, digested by a protease such as trypsin, and analyzed by mass spectrometry (MS)[11]. Collagen peptide mass fingerprinting (PMF) generates a unique set of peptide molecular masses, which can be compared with a database of known sequences to identify the species of origin[12,13]. Tandem mass spectrometry (MS/MS) takes this process a step further by fragmenting the peptide ions in the gas phase to obtain full or partial amino acid sequence, or a set of sequence-specific fragment masses[14,15]. MS/MS offers higher precision in the identification of species than mass fingerprinting and can even provide information about the phylogenetic relationships between different species[16]. PMF and MS/MS are considered to be powerful approaches that are particularly useful for analyzing ancient or degraded samples[5].

A major problem with the current MS-based approaches to species identification is that they are empirical and lack a rigorous statistical framework. Historically, PMF has emerged as the first approach in the early proteomics field for the identification of proteins extracted from gel spots (when the species of origin were well known)[5]. Over the years significant difficulties were uncovered in the PMF approach, and after many attempts to improve the technique, it has finally been abandoned in proteomics due to insufficient statistical robustness [17]. Using the MS/MS approach for protein identification gave improved specificity. For instance, the recent Species by Proteome Investigation (SPIN) technique confirmed the superiority of MS/MS over PMF in terms of species identification quality [18]. Yet the identification of a protein by MS/MS data on a single peptide (“one hit wonder”) is widely considered unreliable [19].

To appreciate the nature of the problem encountered in species identification by MS/MS, it is instructive to review how proteins are currently identified in shotgun (“bottom-up”) proteomics. There, MS/MS data are processed by a “protein search engine” containing a database of all known protein sequences and a decoy database of an equal size, often made of the reversed sequences from the first database. The MS/MS data are then matched by specially designed software simulating MS/MS data from both databases combined, and the matching score between each real and simulated MS/MS spectra is calculated. The score is based on the statistics of matching probabilities, thus depending upon the number of fragments, their type, the peptide length, and the presence of peptides with similar sequences in the database. Sometimes the retention time of peptides[20] and the isotopic distribution of their molecular ions[21] are also utilized for increasing the probability of a correct match. For each peptide, the E-value is then calculated reflecting the expected number of spurious hits with the same or better matching score[22]. Only matches with E-values below an established threshold (typically, 0.05 or 0.01) are accepted[23]. Additionally, false discovery rate (FDR) is calculated based on the number of peptide “hits” from the direct and decoy databases[24].

Subsequently, the peptide matches need to be aggregated into proteins, and this process often contains ambiguity. The parsimony principle is then implemented, demanding peptide attribution to the least possible number of different proteins. Only proteins with at least two unique (i.e., unambiguously attributed) peptides are usually accepted. The FDR of the protein identification is then calculated reflecting the percentage of proteins composed by reversed peptide sequences.

The above procedure is demonstrably complex, but this complexity serves an important purpose – proteomics data are far more trustworthy today than in previous decades. However, still, little of this complexity is implemented in the approaches used for species identification from bone collagen, even though the number of mammal species known (≈6,400 species[25]) is of the same order of magnitude as, e.g., the size of the yeast proteome (≈6000 proteins[26]).

There are also bones of ≈10,000 bird species[27], and ≈32,000 species of fishes[28], which are frequently found in archaeology mixed with other animal bones. Moreover, protein sequences in a proteome are quite diverse, while most collagen proteins from even distant species are highly homologous[29]. This calls for an even more vigorous procedure for species identification by bone collagen analysis than the protein identification routine in regular proteomics. We call this hypothetical procedure “species search engine” or SSE, similar to protein search engines in shotgun proteomics.

Developing a satisfactory SSE will require combined efforts of many bioinformatics groups and will likely take several years. Here we present the first draft of an SSE, named KISSE, based on bone collagen data from 8 species (38 individuals in total): *Halichoerus grypus* (grey seal), *Phoca vitulina* (harbor seal, herein referred to as “seal”), *Ziphius cavirostris* (Cuvier’s beaked whale, “cuvier” for short), *Physeter macrocephalus* (sperm whale), *Eschrichtius robustus* (grey whale), *Falco peregrinus* (peregrine falcon, “falcon”), *Cygnus cygnus* (whooper swan, “swan”) and the extinct *Hydrodamalis gigas* (Steller’s sea cow, “sea cow”). The approach taken does not aim to establish a definitive path for the future but rather focuses on making the most efficient use of current resources. KISSE predicts the most likely species, the next likely species, and estimates the statistical certainty of the results. When the latter value drops below a certain threshold, an indication is obtained that the analyzed species may not be in the database. When a new species is added to KISSE, the scoring and the probability systems need to be recalculated.

Working on the KISSE development, it became necessary to analyze bones from several individuals of the same species for including the species in the database, as the provenance of abone sample is not always known with absolute certainty. As shown in Fig. 1*A*, our initial species collection comprised 5 more species: *Ursus maritimus* (polar bear), *Ursus arctos* (brown bear), *Mirounga leonine* (southern elephant seal), *Balaena mysticetus* (bowhead whale), and *Rangifer tarandus* (reindeer). However, these were later excluded due to the lack of sufficient number of individuals per species (at least three). Bone and tooth samples underwent collagen extraction following Brown protocol[30] (Fig. 1*B*). After collagen extraction, digestion, and desalting procedures, label-free peptides were separated by liquid chromatography and analyzed on a high-resolution mass spectrometer equipped with both Higher Collision Dissociation (HCD) and electron-transfer dissociation (ETD) MS/MS. Proteomics data were recorded in a Data Dependent Acquisition (DDA) mode and the top 5 precursors in each survey mass spectrum were selected for both HCD and ETD MS/MS. This enabled reliable database identification[31] as well as accurate *de novo* sequencing[32], if needed. The MS/MS data were then searched against UniProt/SwissProt reviewed collagenous proteins as well as any protein in the extracellular matrix in the Peaks studio software (Fig. 1*C*).

**Fig. 1.**
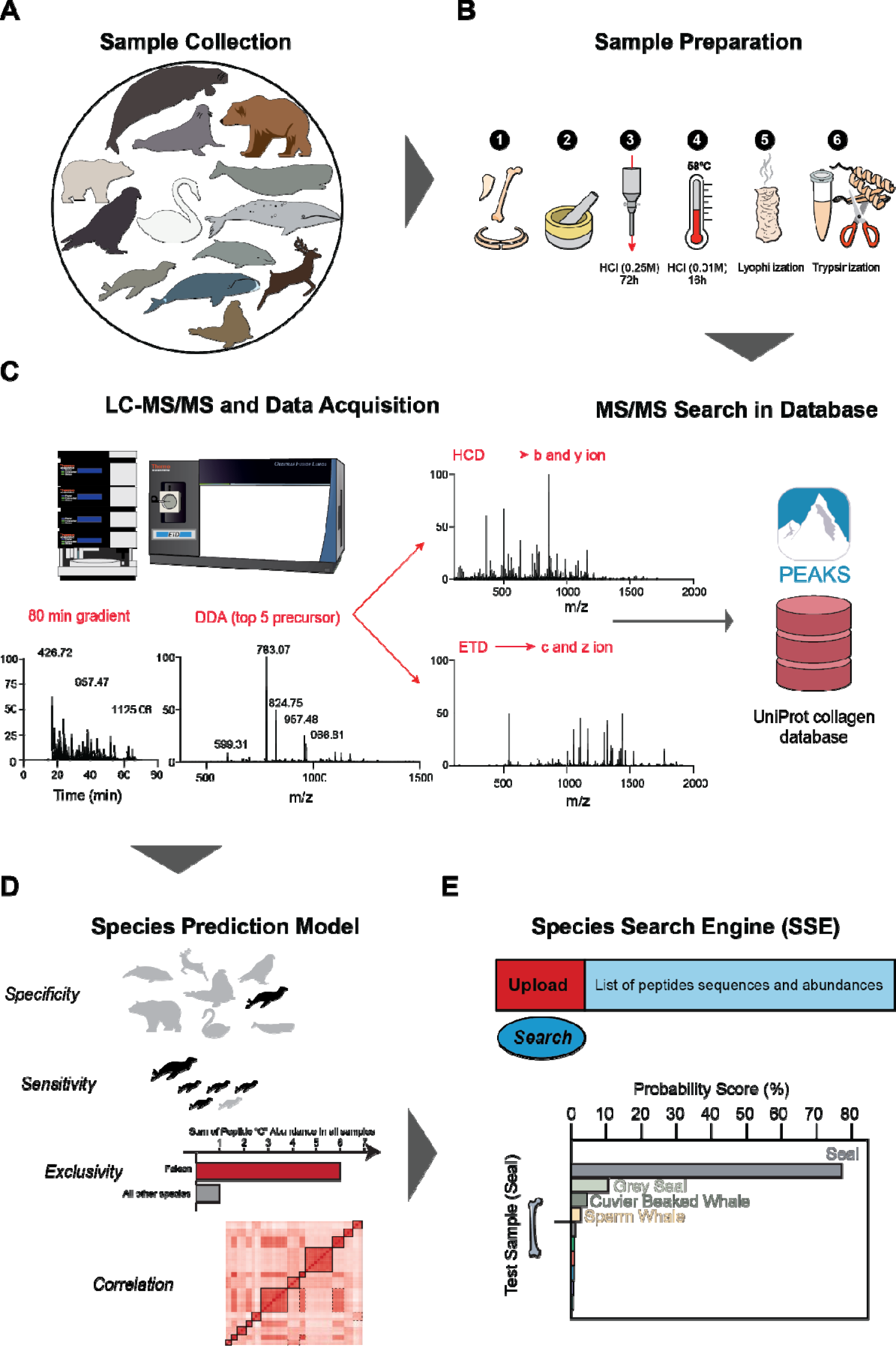
Workflow of developing KISSE – the first Species Search Engine. . *A*, Sample collection. *B,* Collagen extraction from bone pieces: 1. Washing the bone or tooth with water, 2. Grinding the bone or tooth pieces into powder, 3. Demineralization of bone powder with HCl to remove calcium from collagen matrix, 4. Gelatinization with HCl at 58 °C to denature collagen followed by ultra-filtration through a 30 kDa cutoff filter, 5. Lyophilization of denatured collagen solution using a freeze-dryer, 6. Tryptic digestion of collagen. *C*, LC-MS/MS analysis of tryptic peptides, data-dependent acquisition (DDA) with HCD and ETD MS/MS, and usage of PEAKS Studio to perform the search of the .raw files against UniProt collagen database and quantification based on the extracted ion chromatogram of the molecular ions. *D*, Building the Species Predicting model using Specificity, Sensitivity, Exclusivity as parameters for peptide selection, and correlation of peptide abundances as augmenting factor. *E*, User Interface, and the output of the SSE online tool providing the most probable species and the estimated probability score of correct identification.

To build a Species Identification and Prediction by Mass Spectrometry (SIP-MS) model, a subset of peptides was selected based on their specificity, sensitivity, and exclusivity (for details see Experimental Procedures), as well as on the correlation of peptide abundances in individual samples (Fig. 1*D*). The SIP-MS model contains 1) a random forest classification that uses the top N (N=11) most species-specific peptides to predict the taxonomy of a given sample, and 2) a correlation matrix based on the peptide abundances to augment the random forest predictions results and provide the final species identification.

The obtained model for species prediction was then tested based on the out-of-bag (OOB) evaluation technique. To determine the probability score threshold for reliable identification, the model was further tested on known species that were not in the library. The tested model was placed on the web as an R Shiny application with a graphical user interface that provides the most probable species and the estimated probability of correct identification, when the list of peptides and their abundance is uploaded. The external users can also extend the library of specific peptides, adding new species or modifying the existing database (See Experimental Procedure).

## Experimental Procedure

### Sample Collection

Bone and tooth pieces were obtained from the National Museums Scotland, Edinburgh; A.N. Severtsov Institute of Ecology and Evolution, Moscow, Russia; the Swedish Museum of Natural History, Stockholm and Stockholm University, both in Sweden. For more details on individual samples, see Supplementary Table 1.

### Collagen Extraction and Proteomics Sample Preparation

Collagen extraction from bones and teeth was carried out using Brown[30] protocol. At first, each sample was carefully cleaned and washed with deionized water and air-dried. Samples were ground to powder using either a dental drill or by a mortar and pestle. Approximately 50 mg of bone powder from each individual sample were transferred to a glass funnel with an embedded P3 filter (Labglass AB) and glass fiber prefilters (0.7 µm pore size, Merck Millipore) on the top. After the addition of approximately 15 mL of 0.25 M HCl (Merck Millipore) to each glass funnel, samples were incubated at room temperature (RT) for 72 h to remove calcium from the collagen matrix (demineralization/decalcification). At the end of incubation time, HCl was drained from the bottom of the funnel by vacuum, and samples were washed with deionized water twice. To denature the collagenous proteins, 10 mL 0.01 M HCl were added to the glass funnel which was incubated at 58 °C for 16 h. After the incubation, the filtrate solution, obtained from the bottom of the glass funnel using water-jet pump and vacuum chamber, was transferred to Amicon 15 Ultra-filtration 30 kDa centrifugal filter (Merck Millipore) and centrifuged at 4 °C for 20 min at 5000 ×g to remove the contaminants and excess of solvent. The solution remaining on top of the filter was then transferred to a 2 mL Eppendorf tube and kept in the freezer at -20 °C before being lyophilized to obtain sponge-looking dried collagen. Approximately 200 µg of collagen samples were placed at the bottom of each 1.5 mL Eppendorf tube, and 300 µL of ammonium bicarbonate buffer (50 mM in deionized water with pH = 8, Sigma Aldrich) were added to each tube. Samples were vortexed and then incubated at 70 °C for 3 h to solubilize collagen. Following centrifugation at 14,000 ×g for 10 min, the supernatants were collected. Protein concentration was measured using Bicinchoninic acid assay (BCA, Thermo Fisher Scientific). 20 µg of protein from each sample in three replicates were then transferred to a fresh Eppendorf tube and sequence-grade trypsin (Promega) was added to an enzyme: protein ratio of 1: 50. The samples were incubated overnight at 37 °C on a heat block while mixing at 300 rpm. The next day formic acid (FA) (98%, Sigma Aldrich) was added to 5% (v:v, pH in the range of 1 to 3) into each sample to quench the digestion process. Samples were desalted using either C-18 HyperSep™ Filter Plates (Thermo Fisher Scientific) or Sep-Pak Vac C-18 column (Waters) and dried using a Speedvac. Samples were stored at -20 °C before analysis.

### LC-MS/MS Setting and Data Acquisition

Orbitrap Fusion or Lumos mass spectrometers with ETD MS/MS capability equipped with an EASY-Spray source were used online with an UltiMate 3000 RSLC nanoUPLC system (all - Thermo Scientific). The dried samples were dissolved in Buffer A (acetonitrile 2%, FA 0.1%, in water, all from Thermo Fisher Scientific) to a 0.5 µg/µL concentration of the tryptic peptides. The peptide mixtures were preconcentrated before injection using a PepMap C18 nano trap column (length - 2 cm; inner diameter - 75 μm; particle size - 3 μm; pore size - 100 Å; Thermo Fisher Scientific) at a flow rate of 3 μL/min for 5 min. Peptide separation was performed using an EASY-Spray C18 reversed-phase nano-LC column (Acclaim PepMap RSLC; length, 50 cm; inner diameter, 75 μm; particle size, 2 μm; pore size, 100 Å; Thermo Fisher Scientific) at 55 °C and a flow rate of 300 nL/min. Peptides were separated using a binary solvent system consisting of 0.1% (v/v) FA, 2% (v/v) acetonitrile in water (solvent A), and 2% water (v/v), 0.1% (v/v) formic acid in acetonitrile (solvent B). The elution gradient was from 4% B to 15% B for 50 min, to 35% B in 10 min, to 95% B in 3 min, staying at 95% B for 7 min and then decreased to 4% B in 0.5 min and then staying at 4% B for 9.5 min. The mass spectrometry data were acquired in the data-dependent acquisition mode with selection of top 5 precursor ion. In acquiring survey mass spectra, detection was in the orbitrap analyzer with 120,000 resolution in the m/z range 375-1500. The maximum injection time was 50 ms, and Automatic Gain Control (AGC) target was 1e6. MS/MS events comprised two different activation type events, ETD and HCD. In the ETD MS/MS event, the ETD reagent target was 1e6 with the supplemental activation collision energy at 30%. In the HCD MS/MS event, the AGC target was set to 2e5 and the HCD energy at 30%. In both types of MS/MS events the quadrupole filter performed ion isolation with a window of 1.4 m/z, the m/z detection range was set to auto mode and the orbitrap nominal resolution was 50,000. All mass spectra were recorded in profile mode.

### Collagen peptide identification

The MS/MS data were searched against the collagen and extracellular matrix protein sequences in the reviewed UniProt/SwissProt database in PEAKS Studio software version 10.6 (Bioinformatics Solution) using mixed MS/MS activation type (HCD and ETD) and the default setting for all parameters (10 ppm mass error, etc.). Hydroxylation of proline and lysine, deamidation of asparagine and glutamine, and oxidation of methionine were added as variable modifications. The search of MS/MS data against the database was performed with PEAKS DB (with limited number of Post-Transitional Modifications or PTMs), PEAKS PTM (with unlimited number of PTMs), and PEAKS Spider (algorithm that is specially designed to detect peptide mutations and perform cross-species homology search) modules. FDR threshold of 5% was applied to filter the list of identified peptides.

### Peptide quantification

Individual peptide abundances, quantified by measuring the area of extracted ion chromatogram of the molecular ions from PEAKS Studio output, were first normalized by the total peptide abundance in the corresponding sample and then log10 transformed.

### Building the Species Identification and Prediction by Mass Spectrometry (SIP-MS) model

After deciding that at least 3 or more individual samples were needed for each species, the species list was reduced down to 8 groups (Cuvier’s beaked whale, sperm whale, grey whale, grey seal, seal, Steller’s sea cow, peregrine falcon and whooper swan). In total, 26,581 collagen peptides were identified and quantified for these species.

Three parameters were determined for each peptide as follows:

*Specificity: 1/N, where N is the number of species that the given peptide was found in*.

*Sensitivity: Number of individuals from the same species in that a given peptide was found divided by the total number of individual samples for the same species*.

*Exclusivity: Square of the total normalized abundance of a given peptide across all samples of the same species divided by the total normalized peptide abundance across all samples used for model building*.

A subset of 2,469 peptides, here referred to as “informative peptides” was selected. The subset comprised 993 species-specific peptides (those with Specificity and Sensitivity equal to 1) as well as 1476 peptides quantified in at least 25% of all the individual samples with a normalized abundance variation between the species of 3% or higher.

In building the SIP-MS model, a correlation matrix was 1) first calculated for each sample between the abundances of the informative peptides in the sample vs. those in all the other samples. Then these correlations were averaged for all samples from the same species (giving an “averaged correlation” value). A two-sided Student t-test with unpaired and unequal variance between each group was then performed to obtain a p-value that subsequently could be used to obtain a final prediction score. Then 2) in a second step the selection for each species of the most species-specific peptides as a subset of those having Specificity and Sensitivity values equal to 1 were obtained. Since the lowest number (11) of specific peptides was found for the grey seal, for each species the top 11 peptides were chosen after sorting species-specific peptides by the highest Exclusivity.

A random forest model (by “randomForest” package in R version 4.1.0) was then built using the abundances of the selected 88 species-specific peptides (i.e. 8X11 peptides). The model provides the 88 abundances of the corresponding peptides and will give the probabilities (Probability_i_) of a data set to belong to any of the 8 species in the database. In order to deal with missing values (NA values), instead of a constant value, the missing values were replaced by a random numbers X from the range:

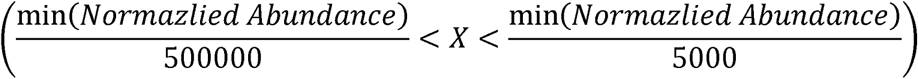

This was a necessary step to ensure that the species identification would be based on the presence of peptides rather than their absence. This correction is important, since in the early attempts for creating the model, peptide abundances were imputed with a small constant value which contributed prominently to erroneous species identification.

The random forest model was trained using approximately two-thirds of the individuals from each species and tested against the remaining one-third of the samples to ensure perfect identification.

For a given sample, the prediction score is finally calculated for each species in the database reflecting the probability, according to the model, of the sample to belong to that species. The Prediction Score_i_ (where *i* is the species index, running currently from 1 to 8) is given by the formula:

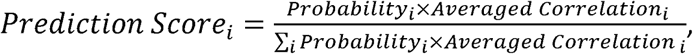

Similarly, the average correlation (Averaged Correlation_i_) is determined from the correlation matrix. Here, the informative peptide abundances for the sample are compared with all individuals in the database, and the correlations are averaged by species.

To identify species, the SIP-MS model requires as input a table with peptide sequences and normalized abundances that needs to be submitted through the graphical user interface. Detailed instructions on how to upload the table are provided on the SIP-MS GitHub page (https://github.com/hassanakthv/SIP-MS). The SIP-MS examines if the informative peptides from the predictive model, and are present in the submitted table and identifies the most likely species. The KISSE then assesses the statistical certainty of species identification.

### Expansion of the SIP-MS model

The SIP-MS model is designed to be expandable, but certain criteria must be met for the submission to the library of new species. Instructions on how and where to submit a new species to the KISSE are provided at the SIP-MS GitHub page (https://github.com/hassanakthv/SIP-MS). Briefly, each sample should be well-documented, preferably with an ID from a museum collection, including information about the bone type, the animal age, sex, place of origin, and other relevant details. For each new species, at least three independent individuals should be available. The added peptide dataset should contain quantified peptide abundances from all individuals. Upon submission the software verifies that the averaged correlation of the peptide abundances between each individual and the rest of the individuals from the same species is higher than 0.7.

## RESULTS

In the database used for building the prediction model, we included 287 proteins identified with at least two unique peptides in bones of 8 species. Of these proteins, 168 were various collagen types, such as Collagen alpha-1(I), Collagen alpha-1(II), Collagen alpha-1(III), etc. On average 158 peptides per collagen type were identified. Of the 26,581 peptides identified in total, 2052 peptides were unmodified, 20,775 were containing proline hydroxylation, 6,480 deamidation of asparagine or glutamine, and 1,079 methionine oxidation. Some of the peptides had several modifications. Among all peptides, 9,582 were unique to specific collagen (57 peptides/per collagen type on average). For the species that did not meet the criteria to be included in training of the prediction model (referred to as "Outside DB"), we identified a total of 248 proteins with at least two unique peptides. In this dataset, 14,483 peptides were included, representing 160 different collagen proteins, with an average of 90 peptides per type. Among these peptides, 917 were unmodified, 11,653 with proline hydroxylation, 3,779 exhibited deamidation of asparagine or glutamine, and 603 carried methionine oxidation. Of these peptides, 4,103 were unique to specific collagens, averaging 26 peptides per type.

### Peptide Selection and SIP-MS Model Building

For each species in our inventory, we discovered collagenous peptides that were unique (Specificity and Sensitivity value of 1) to a given species (Fig. 2). In total there were 993 species-specific peptides while the sperm whale had the highest number of species-specific peptides, with 247 identified peptides common for 3 different individuals. On the contrary, the grey seal gave the lowest number of species-specific peptides, with only 11 common peptides found in 9 individuals. One could hypothesize that with more individuals analyzed, fewer common species-unique peptides will be found, but there was no clear trend between these two numbers. Supplementary Table 2 provides the list of all species-specific peptides along with their protein accession, peptide sequences, mass, charge state, retention time, and averaged normalized peptides abundance.

**Fig. 2.**
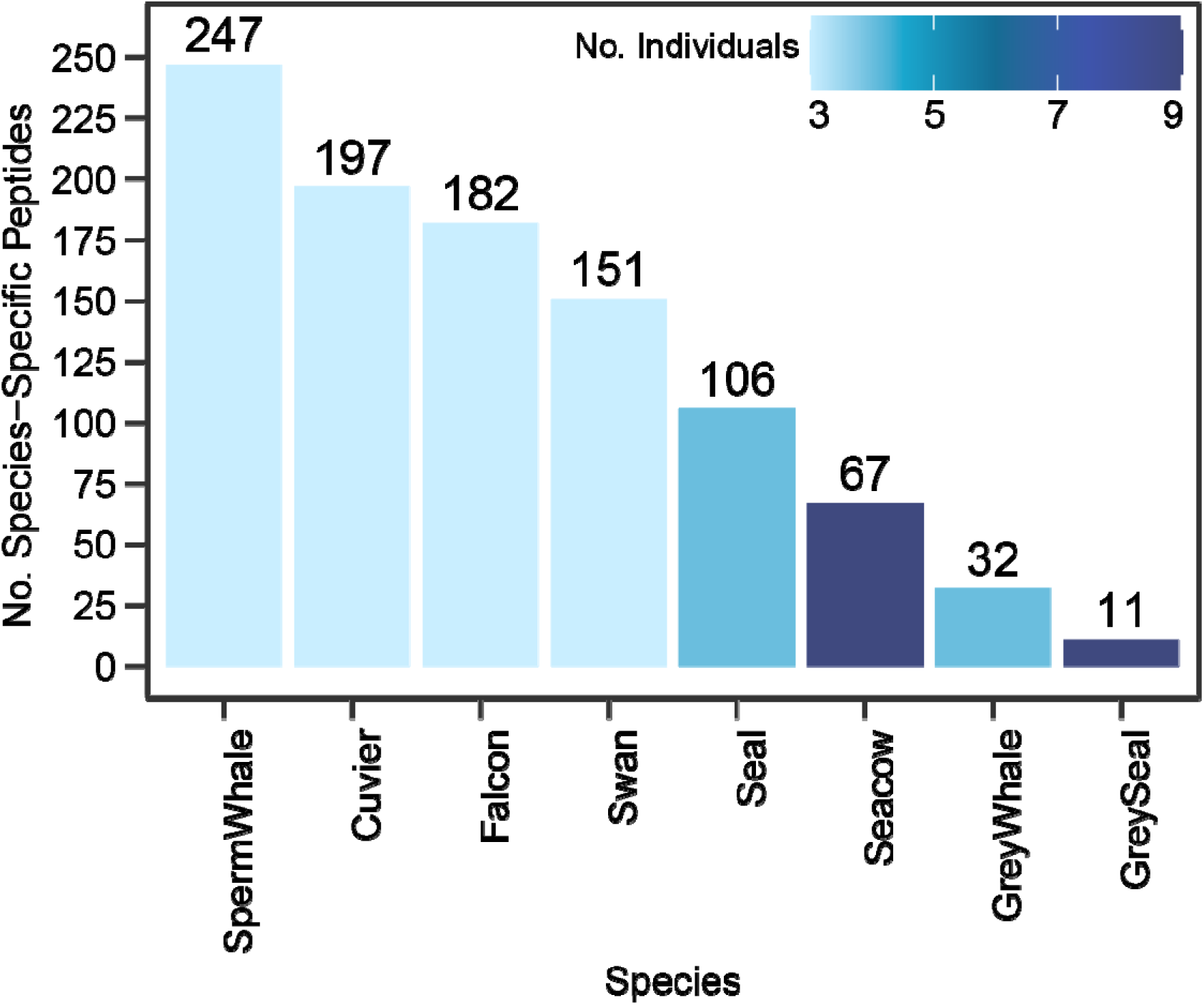
Overview of the Species-Specific peptides. Bar plot for the number of Species-Specific peptides color-coded by the number of individual samples per species.

When the peptide is not present in a mass spectrum, the software assigns to its abundance the noise level. To check the dynamic range of such peptide quantification, we created a heatmap of the normalized abundances for the 88 species-specific peptides selected from the 8 species (Fig 3). Not surprisingly, only clustering of the samples were observed according to the same species, with the dynamic range of abundances stretching for at least 12 orders of magnitude.

**Fig. 3.**
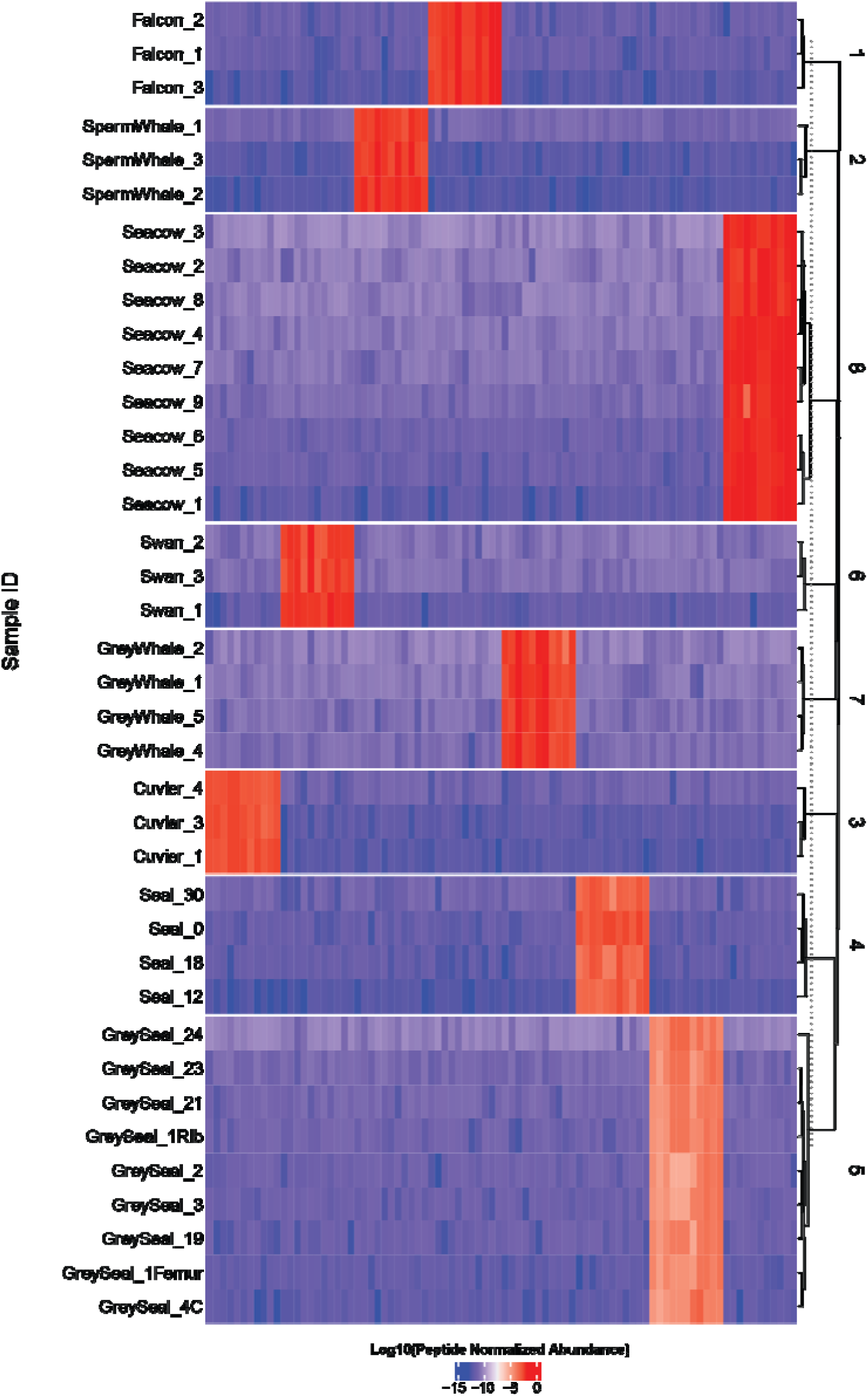
Clustering of 88 species-specific peptides selected for random forest classification in the SIP-MS model.

In total, 2,469 other peptides fitted the criteria for informative peptides (see Experimental Procedure). Abundance correlations for these informative peptides in individual samples are shown in Fig. 4. Not surprisingly, a much higher correlation was observed for individuals from the same species, but a weaker correlation with related species (grey seal and seal, sperm, grey and Cuvier’s whales, falcon and swan, etc.) and absence of correlation for distant species was also observed.

**Fig. 4.**
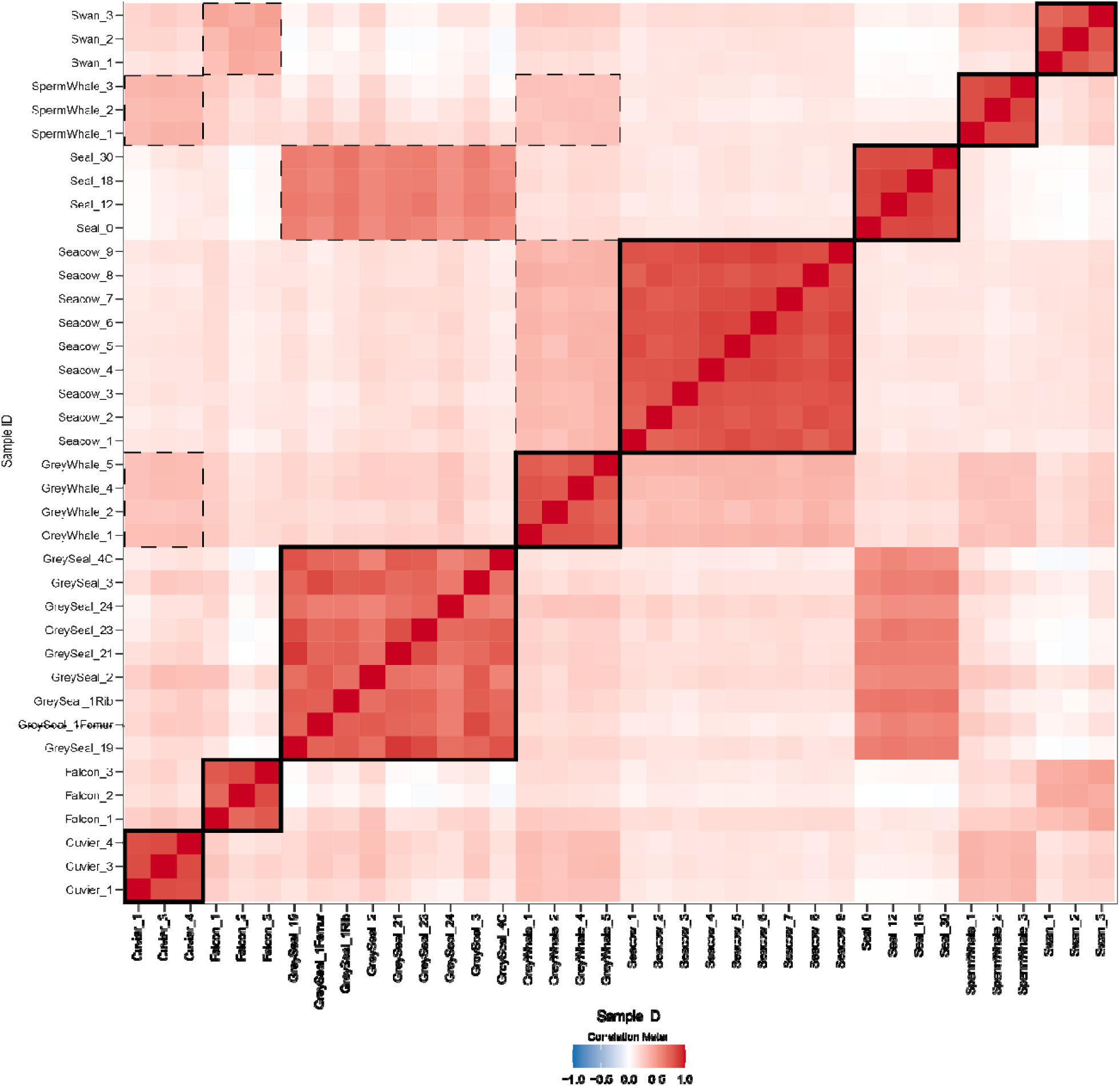
Correlation between the abundances of informative peptides in individual samples. . Thick boxes show the clustering for individual samples within each species group and dashed-line boxes denote notable inter-species correlations.

As expected, the random forest classification SIP-MS model perfectly identified the species included in the training set (Fig. 5*A*). The averaged correlation_i_, which comes from averaging the correlations of the informative peptides of all individuals within each species, is also correctly determining the correct species (Fig. 5*B*). The final prediction scores for each individual sample ranged from 92% to 100% for the correct species identification (Fig. 5*C*). For each species there was a strong significant difference (median p-value < 5×10^-15^) between the prediction of correct ID versus incorrect identification, taking all individuals of every species as a group.

**Fig. 5.**
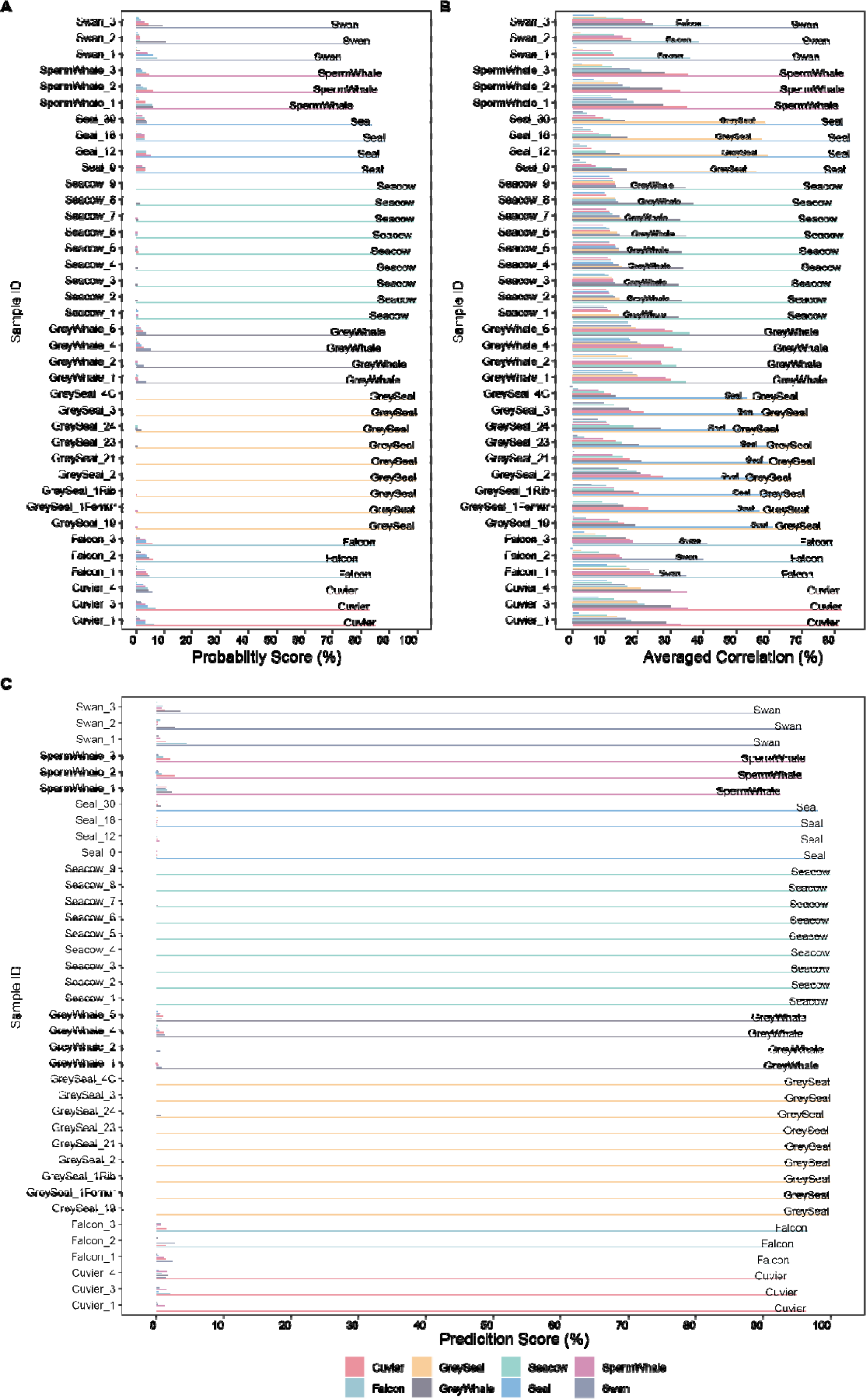
SIP-MS prediction results for species included in the database. *A*, Probability scores from the random forest classifier. *B*, Averaged correlations obtained from informative peptides abundances. *C*, Final Prediction Score from the SIP-MS model.

Thereafter we evaluated how the model performs on the five species that were not used for building the SIP-MS model (“outsideDB” set). For the outsideDB set, peptide abundances were first normalized by the total abundance in each sample, as was done with the training data. To enable comparison with the current database, we retained only the peptides present among the informative peptides in the outside database. The averaged correlation cut-off was set at 0.6, since the least correct correlation species in our dataset (Grey Seal) had the averaged correlation of 0.63. If no species correlates better than the cut-off averaged correlation with the test sample in the SIP-MS model, the KISSE assumes that the species is not in the dataset and uses an averaged correlation to provide the identification of the closest species for that sample.

In Fig. 6, it is shown that none of the OutsideDB samples exhibit an averaged correlation surpassing the correlation cutoff on SIP-MS. Therefore, KISSE concluded that the correct species for these samples are not present in the current database. Instead, KISSE provided the averaged correlation as a means of indicating the most similar species to each sample. In our dataset, the most similar species to bowhead whales is the grey whale, while for brown and polar bears, it is the grey seal (with a median p-value of 10^-7^ for the most similar species).

**Fig. 6.**
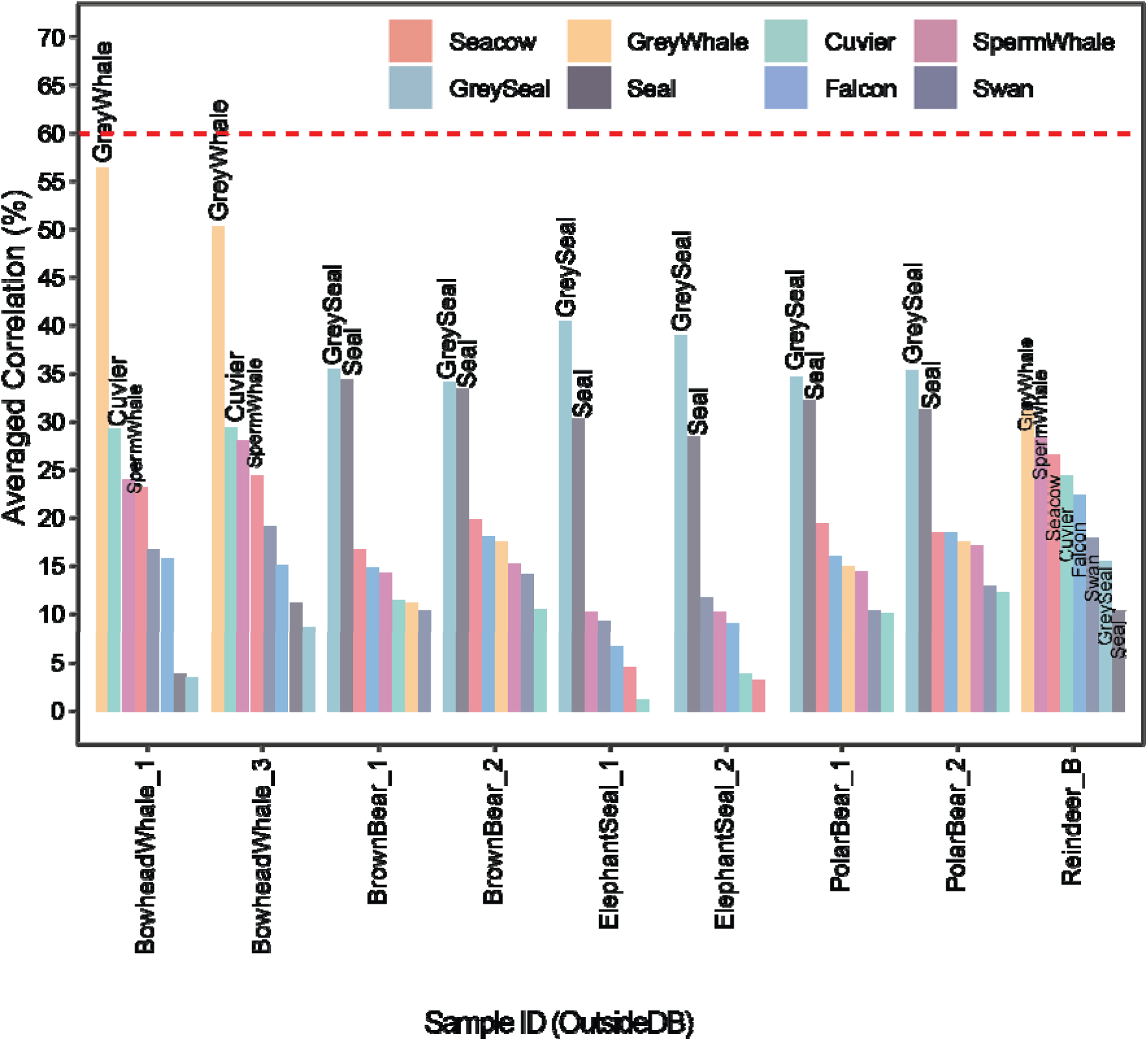
Averaged correlations from the SIP-MS model for samples of species not included in the database, i.e. outsideDB set. The red dashed line shows the averaged correlation cut-off used by KISSE to determine if a species exists in our current database.

Given the absence of terrestrial mammal species in our current database, we did not anticipate a distinctive similarity between reindeer and any of our species. This lack of distinction is evident in the proximity of the averaged correlation and the higher p-value (5×10^-3^), compared to KISSE’s predictions for other OutsideDB samples.

### Comparison of Species-Specific Peptide Identification: MS/MS versus PMF Approaches

To compare the mass sensitivity of species-specific peptide identification in PMF and MS/MS approaches we examined the m/z values of the 993 species-specific peptides founded by MS/MS. We found 35 peptide pairs from two different species that fall within a mass difference of less than 20 ppm, and 43 peptide pairs for the mass difference of less than 25 ppm. As the rule of thumb is that X ppm mass tolerance rules out (100 – X)% of peptides[33] with identical nominal mass, a ≤20 ppm mass tolerance is usually recommended for differentiation of such peptides. Thus, the 35 pairs ambiguous within 20 ppm were excluded from the list of sequence-specific peptides. The resultant number of species-specific peptides identified in different species ranged from 6 (in 9 individual grey seals) to 218 (in 3 individual sperm whales). Since at least 9 species-specific peptides are deemed to be necessary for reliable identification by PMF[34], grey seals would not be reliably identified by the PMF approach.

## Discussion

Here, we create a first installment of an expandable KISSE that can be exploited for the identification of species based on the MS/MS analysis of bone samples using our SIP-MS model. By analyzing bone samples from 8 different species, we determined that a reliable SSE cannot be built based on sequence-specific peptides only. Therefore, we generated a peptide abundance model based on a subset of all quantified peptides.

Although our current SIP-MS model performed perfectly in identifying species for the tested samples, its performance can be further improved with a larger number of individuals within each species. During the model-building stage, we noticed fluctuations in the probability of species identification when splitting the training and test datasets. For example, species with more individuals, such as sea cow and grey seal with 9 individuals, resulted in a higher prediction score (average of 100%) for the most probable species identification compared to species with fewer individuals (average prediction score of 92%).

We should also address the limited number of reviewed annotated proteins available for the species studied in this article. As of November 2024, there are only 76 annotated proteins in SwissProt (reviewed) for these species, and none of them are collagenous proteins. For example, harbor seal (herein referred to as “seal”) has the highest number of annotated proteins (33), while whooper swan and Steller’s sea cow have no annotated proteins at all. Another interesting observation is that the peregrine falcon and whooper swan had a substantial number of species-specific peptides (182 and 151, respectively). Of these, 91 peptides in the falcon and 75 in the swan were annotated as belonging to collagenous proteins of *Gallus gallus* (chicken). Thus, with the available protein databases both these two samples would have been identified as chicken.

The expandable nature of our SIP-MS model allows for future integration of data on other species, enabling the development of a more comprehensive SSE for accurate species identification in fields such as paleontology, archaeology, and forensics.

## Conclusion

The presented SIP-MS model is the very first step toward a universal SSE. It is also a proof-of-principle that even a small sample size (as low as three individuals) can yield reliable species identification in a field where the number of annotated proteins is limited for plenty of species. This approach would greatly benefit from including more collagen-like proteins in the sequence databases.

### Data availability

The mass spectrometry proteomics data have been deposited to the ProteomeXchange Consortium via the PRIDE [35] partner repository with the dataset identifier PXD060919. The KISSE GUI source code is available on GitHub (https://github.com/hassanakthv/SIPMS).

## Acknowledgment

We would like to thank Zena Timmons, Daniela C. Kalthoff, Gunilla Eriksson, and Andrew C. Kitchener for providing bone samples. Additionally, we are grateful to Carina Palmberg and Marie Ståhlberg for their assistance with LC-MS instrumentation.

## Notes

### Competing Interest Statement

The authors have declared no competing interest.

